# A Controlled Test of Risk-Dependent Immune Investment in a Clonal Ant

**DOI:** 10.1101/2025.09.27.678936

**Authors:** Zimai Li, Tze Hann Ng, Jean Keller, Florent Masson, Giselher Grabenweger, Bruno Lemaitre, Nathalie Stroeymeyt, Martin Kaltenpoth, Yuko Ulrich

**Affiliations:** Lise Meitner Group Social Behaviour, Max Planck Institute for Chemical Ecology, Jena, Germany; Faculty of Biological Sciences, Friedrich Schiller University Jena, Jena, Germany; Research Laboratory in Plant Sciences (LRSV), Toulouse University, CNRS, INP, Toulouse, Castanet-Tolosan, France; Department of Insect Symbiosis, Max Planck Institute for Chemical Ecology, Jena, Germany; School of Biological Sciences, University of Bristol, Bristol, United Kingdom; Micropolis, Saint-Léon, France; Department for Plants and Plant Products, Research Group Extension Arable Crops, Agroscope, Zurich, Switzerland; School of Life Science, École Polytechnique Fédérale de Lausanne, Lausanne, Switzerland

## Abstract

Biological systems can benefit from distributing defensive investment unevenly, concentrating protection in parts of the system that face higher risk or hold greater strategic value. In social groups, individuals often face asymmetric infection risks based on their behavioural roles, raising the possibility that immune defences are plastically adjusted according to risk. In social insects, this idea has led to the hypothesis that individuals performing high-risk tasks, such as foraging, should invest more in constitutive immune defences to reduce transmission within the colony. However, testing this hypothesis has been hindered by confounding effects of genotype, age, and infection history. We use the clonal raider ant *Ooceraea biroi* to overcome these limitations. In this system, spontaneous behavioural specialisation between genetically identical, age-matched individuals creates variation in infection risk, while controlling for other sources of variation. We first annotated the *O. biroi* immune gene repertoire and sequenced the transcriptomes of 77 individuals that showed considerable behavioural variation. We then combined fine-scale individual behavioural tracking with three complementary measures of immune investment: immune-related gene expression, antibacterial activity, and survival following infection. Despite behavioural specialisation, we find no evidence that individuals engaging in higher-risk behaviours invest more in constitutive immune defences. These results contradict a long-standing hypothesis in the field and suggest limits to plastic immune allocation based on infection risk in social insect colonies.

## Introduction

All organisms must defend against biological threats such as predators and parasites. However, deploying defensive mechanisms—whether immune, chemical, or behavioural—is typically costly (1,2). Consequently, natural selection is expected to favour strategies that allocate defences efficiently, using them sparingly and where they are most needed.

Across levels of biological organisation, theoretical frameworks predict that defensive investment can be adaptively distributed across parts of a system to maximise protection of the whole (3,4). These frameworks typically assume that optimal defence allocation reflects underlying asymmetries in value or risk: some parts of a system are more valuable or vulnerable than others and are therefore prioritised for protection. For example, at the organismal level, optimal defence theory predicts that plants should concentrate chemical defences in organs of highest fitness value, such as reproductive structures or young leaves, that are critical for survival and reproduction (4). At the population level, epidemiological models predict that targeted vaccination of high-transmission or high-risk individuals can effectively reduce disease spread or severity (5). In both cases, systems benefit when defensive effort is distributed unequally, according to differential threat exposure or strategic value.

Empirical support for this principle is robust at the organismal level (e.g., (6)), where variation in defences across genetically identical structures, such as leaves, can be cleanly attributed to differences in investment. However, testing similar ideas above the organismal level is more difficult. In social groups composed of genetically diverse individuals, variation in immune phenotypes can arise not only from plastic investment in immunity but also from innate genetic differences, or even from social dynamics such as free-riding (7). This makes it challenging to infer whether observed patterns of defence reflect strategic allocation or other factors.

Social insect colonies are promising systems in which to explore this question. They are functionally analogous to multicellular organisms (8), with related individuals cooperating to promote the fitness of the colony as a whole. If strategic immune allocation exists at the group level, it is likely to be observable in social insects. Immune defences in social insects include both individual-level physiological responses and colony-level strategies (9), raising the possibility that immune investment may be plastically adjusted among individuals to optimise colony-level protection, for instance, by reducing pathogen entry or transmission in the colony (10).

In social insects, asymmetries in infection risk are often linked to division of labour. Workers that leave the nest to forage or perform other tasks are generally thought to experience higher exposure to environmental pathogens and are therefore predicted to invest more in constitutive immune defences. Indeed, several studies have reported elevated immune activity in foragers relative to nurses (e.g., (11–14), but see (15) for the reverse pattern). However, these findings are usually confounded by co-variation between behaviour and intrinsic factors such as age or genotype (16), both of which independently affect immune function (17–19). In particular, because foragers are typically older than nurses (20), these patterns are also consistent with ‘inflammaging’, the age-related chronic activation of innate immune defences (21). Moreover, studies based on wild-caught colonies face additional complications: unknown pathogen histories make it difficult to distinguish constitutive immune differences from induced responses to previous or ongoing infections.

To overcome these challenges, we use the clonal raider ant *Ooceraea biroi*, a social insect uniquely suited for disentangling the effects of behaviour from genetic or environmental confounds. Colonies of *O. biroi* naturally consist of genetically identical individuals that emerge in discrete cohorts, enabling the experimental elimination of the usual variation due to genotype and age. As a result, any observed variation in immune phenotype can be attributed to plasticity rather than genetic or age differences. Importantly, genetically identical workers of the same age still establish a stable division of labour in this system: in small experimental colonies, individuals show consistent differences in their tendency to perform tasks inside versus outside the nest (22). This behavioural specialisation in turn creates natural asymmetries in infection risk: previous work has shown that ants working outside the nest are more likely to become infected by parasites than those remaining inside (23).

*O. biroi* thus provides a powerful system to test whether immune investment is plastically allocated among individuals within a social group according to risk. Here, we ask whether individuals facing a greater risk of infection show elevated constitutive immune investment prior to pathogen exposure. Such prophylactic, risk-based variation would constitute a form of “immune division of labour” that could reduce disease transmission and enhance group-level protection. As a first step, we annotate the genomic immune repertoire of *O. biroi* by homology with *Drosophila melanogaster* (24) and compare it to the immune systems of other social insects. We then conduct a fine-scale analysis of individual behaviour in replicate colonies composed of genetically identical, age-matched workers. Finally, we assess three complementary measures of immune investment across individuals: 1) immune-related gene expression, 2) antibacterial activity and 3) survival following pathogen exposure. Contrary to theoretical expectations, we find no evidence for a risk-based prophylactic immune division of labour in *O. biroi* colonies. Individuals that spent more time outside the nests showed an immune investment that was indistinguishable or inferior to that of individuals that stayed in the nest.

## Results

In line with previous studies, *O. biroi* workers in experimental colonies exhibited pronounced behavioural variation despite having identical genotype and age. Our behavioural analysis of eight colonies with ten workers each uncovered individual variation in spatial, temporal and social behaviour. Variation in spatial behaviour, the best-described aspect of division of labour, was evident in extranidal activity (range: 0.004–56.89% of time spent outside the nest; mean ± SD = 10.31 ± 13.44%; Fig. 1A–B; see Methods for definition of behavioural metrics), path length (0.01–271.36 m; 22.95 ± 46.15 m), spatial coverage (2.67–67.33% of grid cells, corresponding to 4–100% of the cells accessible to the ants; 38.18 ± 21.51%), spatial entropy (1.14–6.33; 2.84 ± 1.33) and tortuosity (4.16–38.38; 11.62 ± 5.53). Individuals also varied in their temporal behaviour: some ants moved faster than others (mean velocity: range:1.46–6.84 mm/s; mean ± SD: 2.42 ± 0.91 mm/s), spent a higher proportion of time in movement (locomotor activity: range: 0.004– 37.6%; mean ± SD: 3.95 ± 7.37%), or showed more consistent speed than others (velocity variation: range: 0.77–4.12; mean ± SD: 1.49 ± 0.60); similarly, some ants exhibited bursty nest exits, while others maintained regular intervals (burstiness: −0.11 to 0.79; 0.45 ± 0.15). Finally, some ants had relatively few social contacts on average, while others were nearly always in contact with every other colony member, likely because they stayed in the nest (mean network degree centrality: range: 3.11–8.81; mean ± SD: 7.27 ± 1.43).

**Figure 1.**
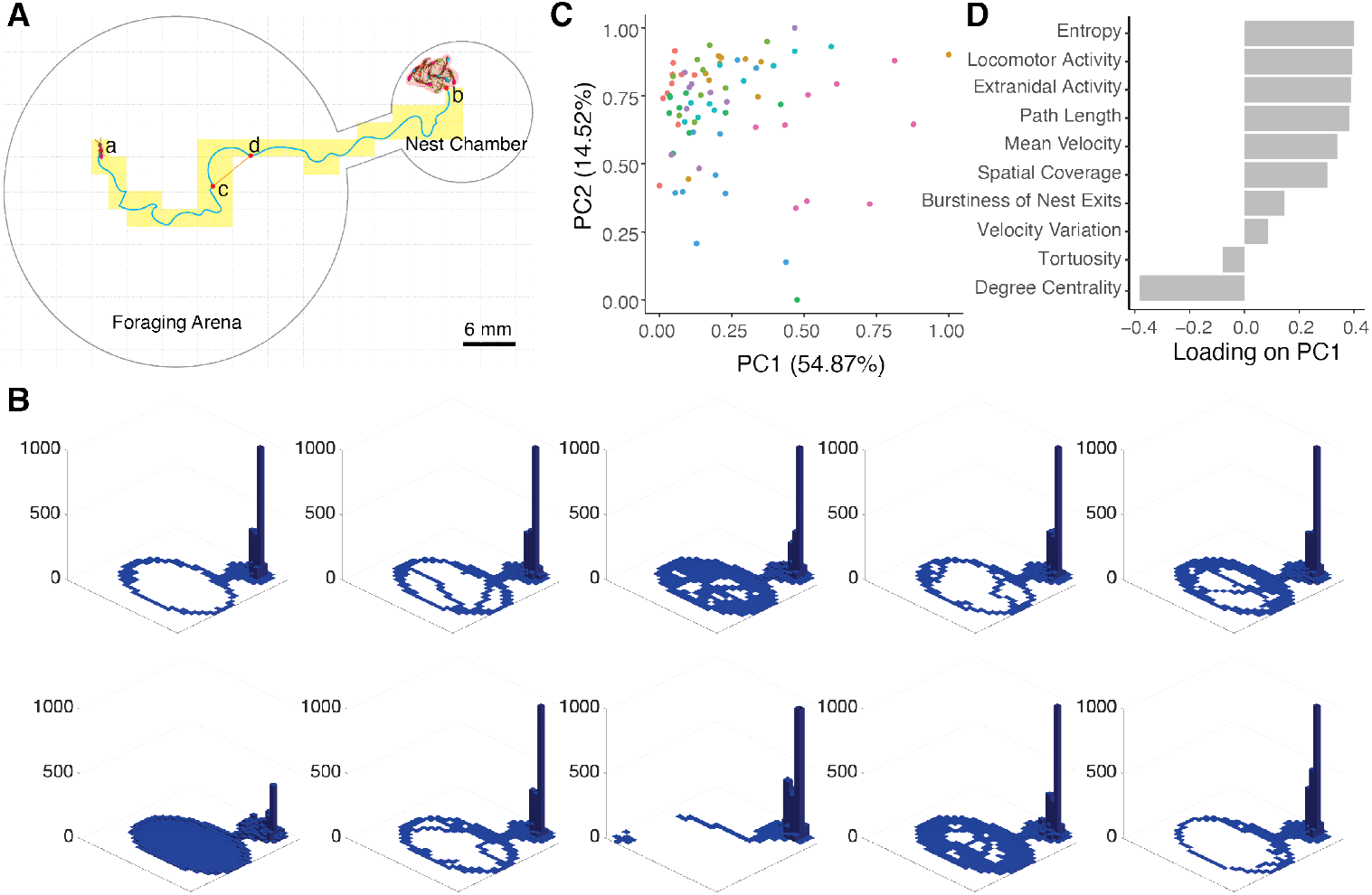
Behavioural variation among genetically identical workers. (A) Schematic representation of behavioural experiment setup and behavioural metric calculation. The blue line represents the trajectory of one ant. The total length of the blue line between points a and b represents path length; the trajectory length (blue line between points c and d) divided by the Euclidean distance (orange line between points c and d) represents tortuosity; the red shaded area in the nest chamber represents the nest; the yellow shaded bins are used to calculate spatial coverage and entropy. (B) Density map of the spatial distribution of 10 genotype- and age-matched ants from one colony. Each subpanel represents one ant. The X and Y axes represent spatial coordinates, and the Z axis indicates the time (in minutes) spent in each grid cell. (C) PC1 and PC2 from the PCA on all behavioural metrics. Each point represents one individual; colours represent colonies (n = 77 ants from 8 colonies). (D) Loadings of individual behavioural metrics on PC1.

PCA revealed no clear clustering of individuals in behavioural space (Fig. 1C), indicating a continuous spectrum of behavioural phenotypes rather than discrete behavioural roles. The first three principal components (PC1–PC3) collectively explained over 80.31% of the total behavioural variance (Supplementary Table S1). PC1 (54.87% of variance) captured spatial exploration (high (>0.3) positive loadings for entropy, extranidal activity, path length, spatial coverage) and general activity level (locomotor activity, mean velocity) while reflecting reduced social interactions (strong (<-0.3) negative loading for degree centrality) (Fig. 1D). Therefore, individuals with high PC1 scores exhibited behavioural profiles classically associated with foragers, with high motility and extranidal activity, and fewer social contacts compared to nurses. PC2 (14.52% variance) predominantly captured temporal consistency (high negative loading for velocity variation), with high scores reflecting stable movement velocity (Supplementary Fig. S1A). PC3 (10.92% variance) captured both spatial behaviour (high positive loadings for tortuosity) and temporal behaviour (high positive loadings for burstiness of nest exits) (Supplementary Fig. S1B).

To investigate the association between individual behaviour and immune-related gene expressions, we first identified 256 immune-related genes in the genome of *O. biroi* (Fig. 2A and Supplementary Tables S2-4). The *O. biroi* genome encodes nearly all key components of the NF-κB pathways IMD, which responds to peptidoglycans (PGN) found in Gram-negative bacteria and some Gram-positive bacilli (Supplementary Table S2), and Toll, which responds to PGN from Gram-positive bacteria and β-glucans from fungi (Supplementary Table S3) (25). In contrast to the black garden ant *L. niger*, which exhibits a reduced number of pattern recognition receptors in the Toll pathway (26), *O. biroi* possesses all major receptors, including GNBP3 for fungal β-glucans and GNBP1 and PGRP-SA for bacterial peptidoglycan. Therefore, the systemic NF-κB immune pathways in *O. biroi* are structurally complete.

**Figure 2.**
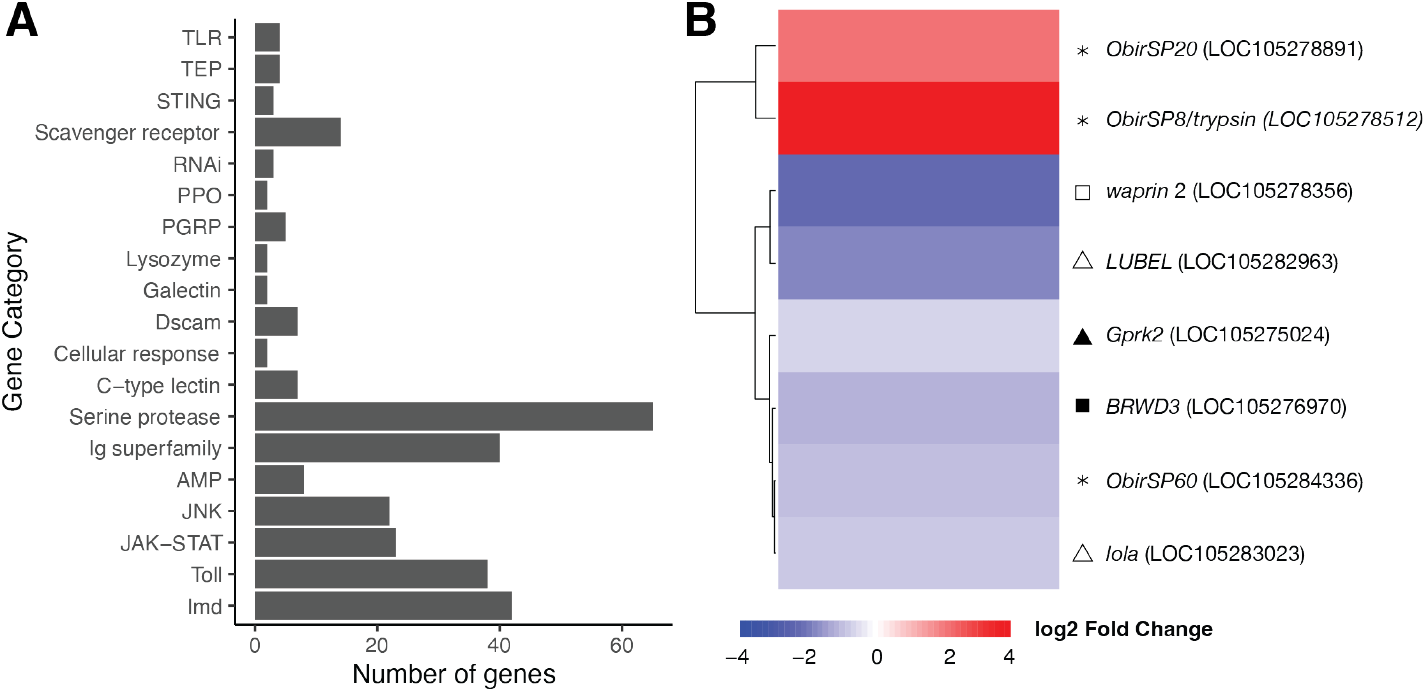
Immune-related gene annotation and differential expression. (A) Repertoire of immune-related genes in *O. biroi*. (B) Eight differentially expressed immune-related genes associated with behavioural PC1. Red indicates upregulation, blue indicates downregulation in more forager-like individuals. The dendrogram represents hierarchical clustering of genes based on expression similarity, using Ward’s D2 method with Euclidean distance. Symbols represent gene categories: ▲ Toll, △ IMD, ■ JAK-STAT, ☐ AMPs, and ✽ serine proteases.

We detected a small set of genes encoding antimicrobial peptides (AMPs) that consists of one *hymenoptaecin*, one *defensin*, and one *crustin* (Fig. 2A and Supplementary Table S4). Additionally, two sequences encoding Waprin, an AMP closely related to Crustins and often associated with venom, were identified. Other potential immune- or venom-related peptides, such as SjAPI-like peptidase inhibitors, cuticle structure proteins with putative antimicrobial activity, and ICK-peptides, were also identified (Supplementary Table S4). Similar to *L. niger*, the number of identified AMP in *O. biroi* is low compared to other ants (27) and much smaller than in *D. melanogaster* and the solitary Hymenopteran *Nasonia vitripennis* (28), suggesting that *O. biroi* individual immunity relies on a reduced set of antimicrobial effectors.

Basal expression of immune-related genes across all 77 surviving workers from eight colonies showed only subtle associations with behavioural variation. Among the 256 annotated immune-related genes, the expression levels of eight genes correlated with PC1 scores (Fig. 2B), which captured division of labour between forager-like and nurse-like individuals (Fig. 1D). Six of these eight genes showed reduced expression levels in ants with higher PC1 scores, i.e., exhibiting more forager-like behaviour. These genes included auxiliary immune regulators *Linear Ubiquitin E3 Ligase* (*LUBEL*, log2FC: −2.05, corresponding to a ca. 4.14-fold decrease between individuals with the highest and lowest PC1 scores) from the IMD pathway, and *Bromodomain and WD Repeat Domain Containing 3* (*BRWD3*, −1.36, ca. 2.6-fold decrease) from the JAK-STAT pathway, an AMP linked to venom (*waprin2*, −2.62, ca. 6.1-fold decrease), and a serine protease (*ObirSP60*, −1.13, ca. 2.2-fold decrease). We also observed small reductions in *longitudinals lacking* (*lola*, −1.00, ca. 2-fold decrease; IMD pathway) and *G protein-coupled receptor kinase 2* (*Gprk2*, −0.66, ca. 1.6-fold decrease; Toll pathway). In contrast, two serine proteases were upregulated in ants with higher score on PC1: *ObirSP8/trypsin1* showed a strong increase (+4.44, ca. 21.7-fold increase), and *ObirSP20* a moderate one (+2.35, ca. 5.1-fold increase) in forager-like ants. We observed no differential expression of genes that are generally considered key markers for systemic immunity in insects. For example, key transcription factors (e.g., *relish, dorsal*), pathogen receptors (e.g., *PGRPs*, Toll receptors), enzymatic effectors such as lysozymes and prophenoloxidases, and classic AMPs (e.g., *defensin, hymenoptaecin*) were stably expressed across behavioural phenotypes. We also found no differential expression of immune-related genes associated with the scores on behavioural PC2 and PC3. In summary, we found modest shifts in the expression levels of a small fraction of immune genes to be associated with variation in forager-like behaviour, but no major, systematic differences in constitutive immune investment. Rather, *O. biroi* appeared to maintain relatively uniform immune-related gene expression across individuals with different behavioural roles in the absence of infection.

As two complementary approaches to quantify immune investment, we conducted antibacterial assays with ant lysates and assessed the survival of ants following exposure to fungal pathogens. For antibacterial assays, we generated two samples per colony (for nine colonies): the half of individuals with the most forager-like behaviour and the half with the most nurse-like behaviour, based on their individual behaviour (i.e., loadings on behavioural PC1). We found no association between behaviour and antibacterial activity: lysates of forager-like and nurse-like ants did not differ in their ability to reduce bacterial growth (paired t-test, t = -1.765, p = 0.1156; Fig. 3A) and bacterial viability (paired t-test, t = -1.0785, p = 0.3123; Fig. 3B). Similarly, we found no association between behaviour and survival following fungal exposure. 93.16% (109 out of 117) of ants died within 12 days following exposure to *Beauveria bassiana*, with time to death ranging from 95.17 to over 300 hours (mean ± SD: 181.36 ± 60.91 hours), consistent with previous work (29). However, none of the PC scores summarising behavioural metrics (PC1, PC2, and PC3) correlated with survival (n = 117 ants from 12 colonies; Cox proportional hazards model, PC1: **χ**2 = 0.42, df = 1, p = 0.52; PC2: **χ**2 = 0.18, df = 1, p = 0.67; PC3: **χ**2 = 0.78, df = 1, p = 0.38), indicating that individual behavioural roles do not predict fungal resistance in *O. biroi* workers.

**Figure 3:**
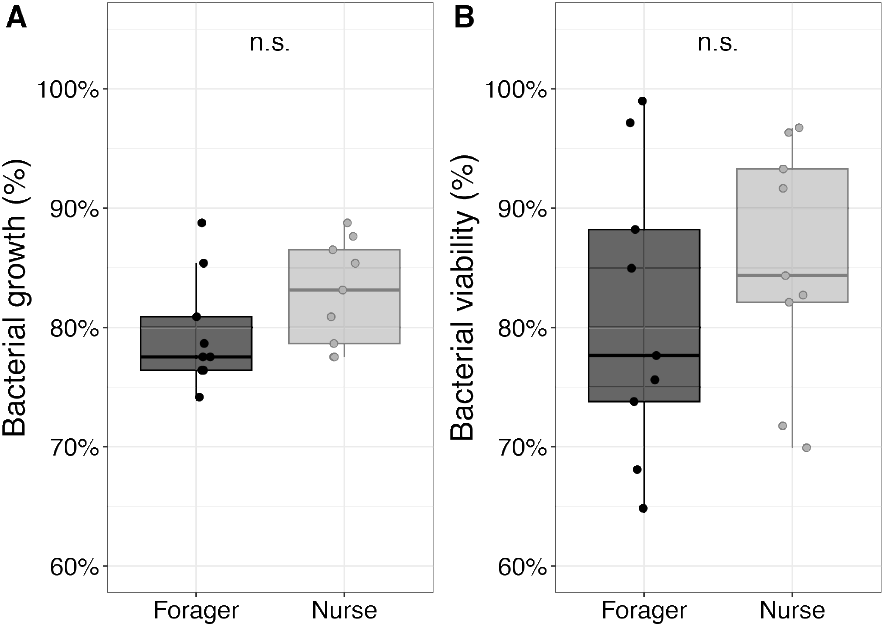
**Antibacterial activity of lysates from forager-like and nurse-like ants**, measured by (A) bacterial growth (OD_600_) and (B) bacterial viability (CFU), relative to negative control. “n.s.”: non-significant.

## Discussion

The unique biology of the clonal raider ant allowed us to investigate associations between division of labour and immune investment in the absence of confounds linked to age, genetics or prior pathogen exposure. Using three measures of immune investment (transcriptomics, antibacterial assays, and survival assays), we found no empirical support for higher immune investment in colony members facing higher infection risk (foragers).

Transcriptomic analyses showed that the basal expression of markers of systemic immune activity in insects, such as pathogen pattern recognition receptors (e.g., PGRPs, Toll receptors), key regulators of canonical NF-κB immune pathways (e.g. *relish, dorsal, spaetzle*), enzymatic effectors (e.g., lysozymes, prophenoloxidase), and classic AMPs (e.g. *defensin, hymenoptaecin*), did not vary across individuals with different behavioural profiles. This contrasts with past work in other social insects, where the expression of some of these genes has been linked to behavioural roles. For example, key immune regulators, including *Toll, Dorsal and Imd*, were expressed 10-to 100-fold more strongly in guards and foragers than in nurses in a stingless bee (14), though this study did not control for age.

Despite extensive inter-individual behavioural variation and large sample sizes, our transcriptomic analysis did not strongly support “immune division of labour”. Among the few (8 out of 256) immune-related genes differentially expressed with respect to individual behaviour, effect sizes and directions varied. In forager-like ants, we observed upregulation of two serine proteases (*ObirSP8/trypsin1* and *ObirSP20*) and downregulation of four secondary immune regulators (*lola, LUBEL, BRWD3*, and *Gprk2*), one serine protease (*ObirSP60*), and one AMP (*waprin*). Serine proteases play roles in prophenoloxidase activation (30) and extracellular Toll signaling activation (31,32). *Gprk2* mediates Toll pathway-driven haemocyte activation (33), *lola* is a non-canonical IMD pathway component (34), *BRWD3* is a positive regulator of the JAK-STAT pathway (35), and *waprin* is a venom AMP with antimicrobial activity (36). As many of these genes have additional non-immune functions (e.g. serine proteases in digestion (37), tissue remodelling (38) and fertilisation (39); *lola* in axon guidance (40)), their expression patterns might be linked to these non-immune functions.

We found no differences in antibacterial activity or in survival upon exposure to pathogenic fungal spores across individuals with different behavioural profiles, so that individual behavioural roles did not predict individual pathogen resistance. Taken together, the absence of modulation in key systemic immune markers, the mixed direction and modest magnitude of effects in a small number of differentially expressed immune-related genes, as well as the lack of association between behaviour and antibacterial activity or pathogen resistance, suggest that immune investment is robust among *O. biroi* individuals that differ in behavioural phenotypes. This contrasts with the majority of previous reports in other social insects, where foragers often showed higher immune activity (e.g., (11–14)). Since pathogen exposure and age were not always controlled in these studies, however, the reported effects could reflect age-related changes in immune function that accompany age polyethism (19,41,42), or differences in past or ongoing infections (43), rather than a prophylactic adjustment of constitutive immune investment to infection risk.

While constitutive defences minimise vulnerability by providing continuous protection, theory predicts that they are not universally favoured by selection (4). This is because mounting and maintaining defences can impose significant fitness costs (44,45). For example, immune investment can reduce resources available for growth or reproduction, generate trade-offs when protection against one pathogen reduces protection against others, or harm the host through immunopathological damage. Therefore, theory predicts that the benefits of constitutive defence will outweigh its costs only when exposure frequency is high, while inducible defences are more advantageous at low exposure frequency (46). Thus, a relatively low exposure frequency, a high cost of maintaining immune defences, or fast and effective induced defences may all contribute to the lack of differences in constitutive immune investment observed here.

Genome annotation revealed that *O. biroi* possesses structurally complete systemic immune pathways, suggesting that it maintains intact individual immune capacity, unlike other ants (26). However, like other social insects, *O. biroi* has a markedly reduced immune gene repertoire compared to solitary insects. For example, we identified only 4 PGRP genes and 5 AMP genes in the *O. biroi* genome, similar to the honeybee *A. mellifera* (4 PGRPs, 7 AMPs), but fewer than in solitary insects like *D. melanogaster* (22 PGRPs, 13 AMPs), *N. vitripennis* (11 PGRPs, 44 AMPs), and *Anopheles gambiae* (7 PGRPs, 11 AMPs) (47–49). Reductions in immune gene repertoires are phylogenetically associated with the evolution of sociality in some (50), but not all (41), social insect lineages, and have been hypothesised to stem from collective immune defences (“social immunity”) reducing the selection pressure on individual immunity (51).

The findings show no consistent evidence that individuals performing higher-risk tasks invest more heavily in constitutive immune defences, with molecular, functional, and survival measures all indicating robust immune investment across colony members. This suggests that immune function may not always be partitioned by task, challenging assumptions in social insect biology and highlighting the potential importance of trade-offs and the buffering effects of social immunity in shaping defence strategies.

## Material and Methods

Three experiments were performed to investigate whether individual behaviour is associated with 1) immune-related gene expression and 2) survival upon pathogen exposure, and 3) antibacterial activity. All colonies within one experiment were processed in parallel, but the different experiments were performed separately.

### Colony setup and behavioural recording

All individuals used in the experiment were reared at 28 °C. To control individual genotypes and age, clonally related workers were sourced from a single age cohort of the same stock colony (genetic lineage A, (52)) on the day they eclosed. The stock colony was free of *Diploscapter* nematodes (23) and any other known ant pathogen. Workers were first housed as a group in an airtight plastic box with a water-saturated plaster of Paris floor for 48 h. Workers were then surface-sterilised following Lacey & Brooks (53) by immersing them in a solution of 70% ethanol for 3 seconds and 1% sodium hypochlorite for 30 seconds, rinsed in autoclaved distilled water, and subsequently housed in sterilised nests, thereby minimising exposure to environmental microorganisms that could induce immune responses. On the day following surface sterilisation, workers were randomly assigned to experimental colonies of 10 individuals housed in air-tight plastic cups (diameter 3 cm) with a floor of autoclaved plaster of Paris saturated with sterilised water. Ants were tagged with unique pairs of paint marks on the thorax and gaster (Uni Paint markers PX-20 and PX-21) for automated tracking. For three weeks, three days per week, plaster floors were cleaned and resaturated with sterilised water, eggs were removed, and colonies were fed with two surface-sterilised brood items of *Messor* minor workers.

To produce larvae for the experiment, a colony was established by separating approximately 1,200 workers from the same stock colony as the experimental colonies. Workers were surface-sterilised and housed in a sterilised nest box, and were maintained as above. Eggs were laid by workers after three days and hatched into larvae after 10 additional days.

When the larvae were five days old, 10 larvae and one experimental group of 10 workers (by then 27-day-old) were placed in recording arenas made of laser-cut cast acrylic. Each arena had a nest chamber (diameter of 15 mm) with autoclaved plaster of Paris floor saturated with sterilised water and a foraging chamber (diameter of 40 mm) of plastic covered with Tyvek^®^ (to make it unattractive as a nesting site), connected by a corridor (2 mm wide and 10 mm long) (Fig. 1A). Workers and larvae were housed in the nest chamber with the corridor closed for 48 h for acclimation and fed once with 2 surface-sterilised brood items of *Messor minor* workers before the start of recording. After acclimation, all food residuals were removed, the corridors were opened, and colonies were filmed continuously for 48 h at 10 Hz and 2,592 × 1,944-pixel resolution, using webcams (Logitech C910).

No food was provided during the experiment. Individuals that died during the observation period were excluded from subsequent analyses.

### Behavioural Data Analysis

The software anTraX (54) was used to extract the raw trajectories of individual ants, i.e., a series of spatial coordinates representing each ant’s position at time intervals of 0.1 s. Raw trajectories underwent preprocessing to identify the brood pile (i.e. the actual nest area within the nest chamber) and missing or aberrant data points, which were then interpolated or removed, respectively, following (55).

Nine behavioural metrics were computed in MATLAB (R2024b) to capture variation in spatial behaviour (extranidal activity, path length, spatial coverage, spatial entropy, tortuosity), temporal behaviour (mean velocity, locomotor activity, velocity variation, burstiness of nest exits) and social behaviour (degree centrality in contact network). These metrics served as proxies for infection risk (56): spatial and social behaviour can affect encounter rates with parasites from the environment or infected social partners, respectively (56–62), while the temporal pattern of movement and contacts can modulate the timing and magnitude of exposure and transmission (63).

Extranidal activity quantified the proportion of time an ant spent outside the nest, and was computed as the amount of time (in video frames) the ant was detected outside the nest (estimated following Jud, Knebel & Ulrich (55)) (Fig. 1A), divided by the total amount of time it was detected. AnTraX assigns the same position to all individuals in the nest, making measures of individual displacement in the nest approximative. Therefore, we only used measures of displacement outside the nest for the calculation of path length, tortuosity, mean velocity, locomotor activity and velocity variation. Path length represented the total distance an ant travelled outside the nest and was computed as the sum of Euclidean distances between consecutive positions at 0.1 s intervals. Displacement exceeding 2 cm s^-1^ were excluded, as such values are biologically implausible and likely reflect tracking errors. To calculate spatial coverage and entropy, a rectangular area encompassing the recording arena (including the foraging arena, the corridor and the nest chamber) was binned into 600 (30 × 20) grid cells, each measuring 2 mm × 2 mm (2mm represents ca. 1 ant body length, Fig. 1A). For each ant, spatial coverage was calculated as the fraction of all grid cells that were visited over the course of the experiment. For each ant, we computed a density map representing the time spent in each cell. Spatial entropy (*H*) was computed from the density map as: *H* = − Σ *p*(*x*)log *p*(*x*), where *p*(*x*) is the proportion of time (in frames) the ant spent in cell *x*. Higher entropy indicates a more even spatial distribution, while lower entropy indicates that the spatial distribution is concentrated in some regions of the arena.

Tortuosity captured the deviation of an ant’s path from the shortest possible route. For each 1-second trajectory segment in which the ant was continuously outside the nest and in movement, tortuosity was defined as the ratio of the actual path length to the net displacement (the Euclidean distance between the start and end points, Fig. 1A). For each ant, overall tortuosity was calculated as the average tortuosity across all trajectory segments. Mean velocity (*V*) was calculated as the average Euclidean distance an ant travelled every second while outside the nest, after excluding values exceeding 2 cm s^-1^, which are biologically implausible and likely reflect tracking errors. Locomotor activity was defined as the proportion of time each ant was moving outside the nest at a velocity exceeding 1 mm s^-1^, after excluding values exceeding 2 cm s^-1^. Workers with high extranidal activity, path length, spatial coverage, spatial entropy, tortuosity, velocity and locomotor activity have a more extensive spatial movement (reflecting forager-like behavioural phenotype), and therefore a higher infection risk (57,58,60) based on past work in *O. biroi*, where foragers had a higher probability of becoming infected by parasitic nematodes compared to nurses (23).

To quantify the temporal variability of each ant’s behaviour, Velocity variation (*VV*) was calculated as the coefficient of variation of the ant’s velocity when outside the nest: 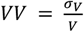, where *σ*_*V*_ is the standard deviation of velocity calculated every second as above, and *V* is the mean velocity. Burstiness of nest exits (B) (from the nest chamber to the foraging arena) was calculated as 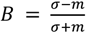where *σ* represents the standard deviation and *m* the mean of time intervals between consecutive nest exits. A burstiness value of –1 indicates a regular pattern of nest exits, a value of 1 indicates clustered (bursty) exits, and a value of 0 indicates that the nest exit followed a Poisson distribution. High velocity variation and burstiness of nest exits indicate irregular activity, which can affect how and when individuals are exposed to environmental pathogens (64). Degree Centrality quantified the average number of contacts each ant had with other ants per frame. It was computed as the total number of contacts across all frames divided by the total number of frames in which the ant was detected. A contact was defined as an event when two ants’ positions were within 2 mm (ca. 1 body length) of each other outside the nest, or when they were both in the nest at the same time. Degree centrality reflects an ant’s risk of becoming infected or transmitting an infection through contact (61,62).

To obtain summary metrics for behavioural profiles accounting for the fact that some behavioural metrics (e.g., extranidal activity and spatial entropy) are expected to covary, we reduced the dimensionality of the behavioural data using Principal Component Analysis (PCA) using the function *prcomp* in R 4.4.2 (65). Missing data (0.65 %) were imputed using the function *imputePCA* from package *missMDA* (66). This method estimates missing values based on the PCA structure of the complete data, ensuring that the overall variance is preserved. Following the Kaiser-Guttman criterion (67), which retains only principal components with eigenvalues greater than 1 (an eigenvalue of 1 indicates that a principal component explains as much variance as an original variable), the first three principal components (henceforth PC1, PC2, and PC3) were retained.

These components cumulatively accounted for 80.31% of the total inter-individual behavioural variation (Supplementary Table S1) and were used in subsequent differential gene expression analysis. The behavioural metric loadings on PC1 were qualitatively similar across experiments, whereas those on PC2 and PC3 differed (Fig. 1D, Supplementary Fig. S1 and S2).

### RNA extraction, library preparation and sequencing

Immediately after the end of recording, 78 surviving workers (out of 80) from eight colonies were flash-frozen in liquid nitrogen and stored individually at −80 °C. RNA from each individual (whole-body samples) was extracted and sequenced by Novogene Co., Ltd (Cambridge, UK). For RNA extraction, individual ants were homogenised in TRIzol and incubated at room temperature (RT) for 5 min, followed by the addition of 0.2 mL of chloroform per 1 mL of TRIzol. Samples were shaken vigorously for 15 s, incubated for another 3 min at RT, and then centrifuged at 12,000 rpm and 4 °C for 15 min. The upper aqueous layer (approximately 0.6 mL) was transferred to a new tube, and RNA was precipitated by adding 0.5 mL of isopropyl alcohol. Samples were incubated at RT for 10 min and centrifuged at 12,000 rpm and 4 °C for 10 min. The RNA pellet was washed once with 75% ethanol (1 mL per mL of TRIzol), vortexed, and centrifuged at 7,500 rpm and 4 °C for 5 min. After removing the supernatant, the pellet was air-dried for 5-10 min at RT and then resuspended in RNase-free water. Dissolved RNA was incubated for 10 min at 55-60 °C to ensure complete solubilisation.

Messenger RNA (mRNA) was purified from the total RNA using poly-T oligo-attached magnetic beads for library preparation. The mRNA was then fragmented, and first-strand cDNA synthesis was performed using random hexamer primers, followed by second-strand cDNA synthesis. The library was constructed through a series of steps, including end repair, A-tailing, adapter ligation, size selection, amplification, and purification. Library quality was assessed using a Qubit fluorometer and real-time PCR for quantification, as well as a Bioanalyzer for size distribution. One sample was discarded due to low quality so that 77 samples were sequenced (150 bp, paired-end) on an Illumina NovaSeq 6000 platform, yielding 44.95 ± 3.86 million reads per sample.

### Bioinformatics

#### Immune-related gene annotation

We annotated immune-related genes in *O. biroi* by drawing on several established gene sets from *D. melanogaster*, the honeybee (*Apis mellifera*) and pertinent literature sources. First, we constructed a query database of 142 predicted polypeptides from *D. melanogaster* genes associated with the NF-κB pathways IMD and Toll, and JAK-STAT and JNK pathways, following the approach of Masson et al. (26). We performed BLASTp (68) searches in Geneious 2021.2.2 (https://www.geneious.com) (BLOSUM62, word size = 3, gap open penalty = 11 and extension = 1) using the *O. biroi* proteome predicted from the Obir_v5.4 reference genome (Accession number GCF_003672135.1; BioProject PRJNA420369). A homologue was accepted upon reciprocal best BLASTp hits (e-value < 10^−20^).

We took additional steps to ensure the accurate identification of homologues of genes belonging to large, highly homologous families: TLRs, PGRPs, TEPs, serine proteases, and C-type lectins. For TLRs and TEPs, candidate sequences identified by reciprocal BLASTp with *D. melanogaster* sequences were blasted a second time against the *O. biroi* predicted proteome to identify distant homologues that may have not been identified by direct alignment with *D. melanogaster* queries. For PGRPs, all known isoforms of *D. melanogaster* PGRPs were queries and *O. biroi* hits were directly validated only if a single reciprocal best match was found. PGRP sequences that failed to show clear reciprocal hits or mapped ambiguously to multiple IDs were subsequently validated using orthology inference with SHOOT. To identify serine proteases, we employed a multi-step process due to high sequence similarity. After an initial BLASTp search using *D. melanogaster* queries, we blasted all hits against the *O. biroi* proteome to identify more distant homologues that may have not been identified by direct alignment with *D. melanogaster* queries. Duplicates were manually removed, and the remaining candidates were scanned on InterProScan to confirm the presence of a peptidase domain. BLASTp candidates with no functional peptidase domain were manually removed, yielding a definitive list of 65 unique serine protease candidates. C-type lectin candidates, drawn from *A. mellifera* annotations (47), were identified using the same reciprocal BLAST method and further annotated based on alignments with *A. mellifera* proteins using MAFFT L-INS-I (69) and phylogenetic analysis (Neighbour-Joining, Jukes-Cantor).

In addition to the systemic immune pathways, we also screened stimulator of interferon genes (STING), Nimrod family proteins (EGF-domain–containing), scavenger receptors, Down syndrome cell adhesion molecule (Dscam; although the role of this gene in phagocytosis has been questioned by recent studies (24)), and Immunoglobulin (Ig) superfamily genes. STING candidates were obtained from Flybase Pathway Reports (70) and validated by best reciprocal BLASTp between *D. melanogaster* and *O. biroi* proteins. Nimrod genes were identified by querying *D. melanogaster* Nimrod polypeptides, pooling hits, and verifiying them reciprocally. Based on Melcarne (71), only 4 were identified to be involved in immunity in Drosophila. Scavenger receptors were obtained from an initial *D. melanogaster* list and blasted against *O. biroi* proteome. After removal of duplicates and manual curation, 10 were confirmed as scavenger receptors, with two showing direct functional links to immunity based on known literature and domain analyses. For Dscam, *D. melanogaster* Dscam1 yielded multiple hits, which were filtered by reciprocal matching and domain checks. Because Ig superfamily annotations are limited in *D. melanogaster*, we referred to *A. mellifera* genome annotation (72) and performed BLASTp plus domain validation in InterProScan (73) to confirm Ig family domains in *O. biroi*, yielding 40 sequences.

Last, we searched for AMPs by appending 3,167 AMP sequences from the APD3 database (2020 release) (74) to our existing query set. BLASTp searches with an e-value <10−5 produced candidate AMPs, which we then blasted against the nr database to eliminate false positives, and scanned on InterProScan (73) to verify the presence of AMP-specific domains. We also searched for other potential immune-related genes, such as prophenoloxidase, galectins, and various cellular immunity markers, using *D. melanogaster* references and reciprocal BLASTp. All identified sequences were compiled and gene IDs were sourced to produce a comprehensive list of 256 candidate immune-related genes in *O. biroi* (Supplementary Tables S2-4).

#### Differential gene expression (DGE) analysis

The transcriptome reads were mapped to the *O. biroi* reference genome (v5.4, BioProject: PRJNA420369) using the nf-core/rnaseq pipeline (75). Raw reads from all samples were trimmed using TrimGalore (76) using a minimum read length of 20 base pairs and a minimum quality score of 30 (an enhancement from the default score of 10). Afterwards, SortMeRNA (77) was implemented to filter and remove ribosomal RNA (rRNA) sequences from the data, ensuring that only mRNA reads were considered in subsequent steps. The filtered reads were then aligned to the reference genome using STAR (default parameters). Afterwards, read counts for each gene were obtained using Salmon (78).

We tested for immune division of labour with a DGE analyis on annotated immune-related genes using the *limma* package (79) in R 4.4.2 (65). The absence of clear clustering in the PCA on behavioural metrics suggested a continuous spectrum of behavioural variation rather than discrete behavioural roles. We therefore used the summary metrics (PC1, PC2, and PC3 scores) as continuous variables in the DGE analysis. Genes with counts per million < 1 in more than half of the samples were excluded prior to normalisation. Trimmed Mean of M-values normalisation was applied, followed by voom transformation. We modelled each gene’s expression as a function of PC1, PC2, and PC3, while accounting for the random effect of colony using the function *duplicateCorrelation* in *limma* (80). Significance was determined at a False Discovery Rate (FDR) ≤ 0.05 using Benjamini–Hochberg–adjusted p-values to correct for multiple testing in each DGE analysis.

Because behavioural data were collected continuously over a 48-hour period, but RNA was sampled at a single time point at the end of this period, there was a temporal mismatch between the behavioural and the transcriptomic data. To assess whether this mismatch influenced our DGE results, we repeated the analysis using behavioural metrics calculated only from the final 12 hours of the tracking period. 84.55% of behaviour-associated DEGs in the final 12 hours of tracking were also associated with behaviour in the entire 48-hour period of tracking (Supplementary Fig. S3), suggesting the association between behaviour and gene expression was robust over the duration of the experiment. We present the results of the DGE analysis based on the full experiment (48-hour) here.

We performed gene set enrichment analysis using Gene Ontology (GO) annotations following Li et al. (23). Enrichment was tested with the R package *topGO* (81), applying the *elim* algorithm to account for GO topology. GO terms with p-values < 0.05 were considered significantly enriched. To aid interpretation, enriched terms were semantically clustered using the package *rrvgo*.

### Fungal exposure assay

Following behavioural recording, we measured the survival of ants from 12 experimental colonies after exposure to the generalist entomopathogenic fungus *Beauveria bassiana* isolate ART2850 (isolated from a *Technomyrmex vitiensis* ant in Zurich, Switzerland, and stored in the stock collection of Agroscope, Zurich, Switzerland). A suspension of 10^8^ conidiospores mL^−1^ in sterile 0.05% Triton X-100 was prepared, and its germination rate was verified to exceed 95% on Sabouraud dextrose agar at 23 °C for 20 h. Ants were individually immersed in a separate 100 μL of the suspension for 5 seconds, placed on filter paper for 10 min to dry, and then transferred into an air-tight plastic cup with an autoclaved plaster of Paris floor saturated with autoclaved water. In addition to the ants from experimental colonies, 10 workers from the stock colony underwent the same procedure but with sterile 0.05% Triton X-100 lacking conidiospores as controls. Survival was monitored daily for 13 days post-exposure. On the day of death, each cadaver was surface-sterilised using the same method as for live ants, placed in a Petri dish with damp filter paper, and observed for 14 additional days to track any fungal outgrowth. Only ants with confirmed fungal outgrowth on their cadavers were included in survival analyses.

To test the relationship between behaviour and resistance to a fungal pathogen, we conducted a survival analysis using a Cox Proportional Hazards Model (implemented using the *coxph* function from the R package *survival*). The model included individual scores on the first three behavioural principal components (PC1, PC2, and PC3) as continuous predictors and colony identity as a random factor to account for colony-level variation. We initially fit a model that included all two-way interactions among the three principal components, none of which were statistically significant. The final model therefore included only the three principal components as predictors. The proportional hazards assumption was verified using standard diagnostic plots and Schoenfeld residual tests (*cox*.*zph* function in the *survival* package). Statistical significance was assessed using likelihood ratio tests (LRT).

### Antibacterial activity

Following behavioural recording, 84 workers from nine colonies were used in an *in vitro* antibacterial assay modified from (82). Ants were flash-frozen in liquid nitrogen and stored individually in 1.5 mL Eppendorf tubes at -80 °C. Total protein was extracted from individual frozen whole-body samples homogenised in 100 µL ice-cold phosphate-buffered saline (PBS; Sigma) with 1 mM Phenylthiourea (PTU; Sigma). The lysate was immediately centrifuged at maximum speed for 10 min at 4 ºC, and the supernatant was transferred to a new tube. The supernatant was centrifuged again to remove any debris, and the supernatant was transferred to a new tube. Because antibacterial activity could not be measured from single ants, we pooled supernatant from the most forager-like ants and the most nurse-like ants in each colony (3 to 5 individuals per sample, depending on ant survival during recording), based on loadings on behavioural PC1, for a total of 9 samples per behavioural group. The protein concentration of each sample was determined using the Bradford assay and samples were frozen at -80 ºC for further use.

*Escherichia coli* TOP10 (pUC18) was cultured in Luria-Bouillon (LB; Sigma) broth with carbenicillin (200 µg/mL; Carl Roth) at 37 ºC overnight. The overnight *E. coli* culture was pelleted at 3000 rpm for 10 min and prepared in sterile PBS at OD_600_ = 1. Samples (1 µg of total protein) were mixed with 25 µL of bacteria suspension adjusted to 5 × 10^6^ CFU/mL in LB with carbenicillin. Reactions in 1.5 mL tubes were incubated for 3 h at 37 ºC, then each reaction was diluted with 190 µL LB with carbenicillin and 100 µL were immediately transferred to 96-well plate in duplicates for bacterial growth determination by OD_600_ reading. The remaining 10 µL were further diluted and plated on LB plates with carbenicillin in duplicates and incubated overnight at 37 ºC for bacterial viability determination by colony-forming units (CFU) count. OD_600_ readings and CFU counts for each behavioural group were normalised to the negative control (PBS without lysate) to calculate relative bacterial growth and viability. Values were averaged across technical duplicates and compared between forager-like and nurse-like samples using paired t-tests.

## Acknowledgements

The authors thank Fionna Knecht for training, Daniel Veit for help building the experimental setup, Daniel Knebel for help setting up experiments, Martin Niebergall for help establishing the computational environment, and Heiko Vogel for early advice on bioinformatics. This work was supported by the Max Planck Society and the European Research Council (ERC) under the European Union’s Horizon 2020 research and innovation programme (grant agreement no. 851523) to Y.U.; NS and FM acknowledge funding by the European Research Council (ERC Starting Grant ‘DISEASE’, no. 802628, to NS). This is Clonal Raider Ant Paper # 41.

## Author contributions

Z. L.: conceptualization, data curation, formal analysis, investigation, methodology, validation, visualization, writing—original draft, writing— review and editing; TH. N.: investigation, methodology, visualization, writing— original draft, review and editing; J. K.: methodology, writing— review and editing; F. M.: formal analysis, writing— review and editing; G. G.: resources, writing— review and editing; B. L.: resources, writing— review and editing; N. S.: resources, writing— review and editing; M. K.: resources, writing— review and editing; Y.U.: conceptualization, funding acquisition, methodology, project administration, resources, supervision, writing—review and editing.

