## Supplementary Table for "A Controlled Test of Risk-Dependent Immune Investment in a Clonal Ant"

**Table S1.** Summary of behavioural principal component analysis (PCA) in the experiment investigating the association between individual behaviour and immune-related gene expression.

| Dimension | Eigenvalue | Variance (%) | Cumulative Variance (%) |
|---|---|---|---|
| 1 | 5.49 | 54.87 | 54.87 |
| 2 | 1.45 | 14.52 | 69.39 |
| 3 | 1.09 | 10.92 | 80.31 |
| 4 | 0.77 | 7.66 | 87.97 |
| 5 | 0.53 | 5.28 | 93.25 |
| 6 | 0.41 | 4.07 | 97.33 |
| 7 | 0.16 | 1.56 | 98.89 |
| 8 | 0.06 | 0.61 | 99.50 |
| 9 | 0.04 | 0.44 | 99.95 |
| 10 | 0.01 | 0.05 | 100.00 |

**Table S2.** *O. biroi* IMD pathway gene list.

| | Symbol | Flybase ID | O.biroi Homologue CDS ID | O.biroi Homologue Gene ID | BLAST e-value vs D. melanogaster homologue |
|---|---|---|---|---|---|
| Key Components | Imd | FBgn0013983 | RLU16836.1 | LOC105275888 | 6.92E-17 |
| | Fadd | FBgn0038928 | RLU18442.1 | LOC105281606 | 2.33E-13 |
| | PGRP-LC | FBgn0035976 | RLU18495.1 | LOC105279031 | 2.89E-60 |
| | Tak1 | FBgn0026323 | RLU18624.1 | LOC105284644 | 5.72E-108 |
| | Diap2 | FBgn0015247 | RLU20359.1 | LOC105282023 | 1.51E-97 |
| | Rel | FBgn0014018 | RLU20433.1 | LOC105282066 | 1.26E-94 |
| | IKKbeta | FBgn0024222 | RLU24379.1 | LOC105277074 | 6.30E-44 |
| | Dredd | FBgn0020381 | RLU27033.1 | LOC105280305 | 1.28E-30 |
| | Tab2 | FBgn0086358 | RLU27115.1 | LOC105283447 | 5.66E-14 |
| Secondary Components | lola | FBgn0283521 | RLU14810.1 | LOC105283023 | 1.05E-69 |
| | Ubc4 | FBgn0015321 | RLU15572.1 | LOC105287045 | 1.67E-93 |
| | Parp | FBgn0010247 | RLU15812.1 | LOC105277761 | 0 |
| | SkpA | FBgn0025637 | RLU16333.1 | LOC105280685 | 2.24E-90 |

| | | | | |
|---|---|---|---|---|
| LUBEL | FBgn0031857 | RLU16718.1 | LOC105282963 | 4.25E-14 |
| lwr | FBgn0010602 | RLU17305.1 | LOC105285009 | 1.41E-96 |
| sick | FBgn0263873 | RLU17428.1 | LOC105282273 | 0 |
| kay | FBgn0001297 | RLU17728.1 | LOC105286045 | 2.16E-21 |
| PGRP-SC2 | FBgn0043575 | RLU17731.1 | LOC105283691 | 1.87E-51 |
| PGRP-LB | FBgn0037906 | RLU17733.1 | LOC105283690 | 9.78E-48 |
| Uev1A | FBgn0035601 | RLU18291.1 | LOC105276007 | 2.54E-83 |
| flfl | FBgn0024555 | RLU18980.1 | LOC105283106 | 0 |
| akirin | FBgn0082598 | RLU19462.1 | LOC105275525 | 2.99E-42 |
| Pp4-19C | FBgn0023177 | RLU19546.1 | LOC105275704 | 0 |
| Su(var)2-10 | FBgn0003612 | RLU19689.1 | LOC105287871 | 3.16E-159 |
| UbcE2M | FBgn0035853 | RLU21266.1 | LOC105284092 | 1.56E-106 |
| PPP4R2r | FBgn0030208 | RLU21272.1 | LOC105276511 | 3.60E-54 |
| scny | FBgn0260936 | RLU21301.1 | LOC105284507 | 2.65E-158 |
| eff | FBgn0011217 | RLU22441.1 | LOC105287597 | 6.01E-110 |
| Cul1 | FBgn0015509 | RLU23105.1 | LOC105287023 | 0 |
| dnr1 | FBgn0260866 | RLU24327.1 | LOC105277436 | 1.00E-66 |
| velo | FBgn0035713 | RLU24822.1 | LOC105277202 | 4.54E-128 |
| Drice | FBgn0019972 | RLU24837.1 | LOC105278081 | 1.02E-120 |
| casp | FBgn0034068 | RLU25369.1 | LOC105278158 | 4.32E-156 |
| Traf6 | FBgn0265464 | RLU25883.1 | LOC105280100 | 4.47E-72 |
| trbd | FBgn0037734 | RLU26285.1 | LOC105284629 | 0 |
| puc | FBgn0243512 | RLU15251.1 | LOC105280616 | 5.17E-70 |
| PGRP-SD | FBgn0035806 | RLU16632.1 | LOC105274606 | 7.71E-48 |
| CYLD | FBgn0032210 | RLU18404.1 | LOC105276204 | 4.30E-96 |
| ben | FBgn0000173 | RLU20330.1 | LOC105282035 | 2.27E-102 |
| bsk | FBgn0000229 | RLU21971.1 | LOC105285161 | 0 |
| His2Av | FBgn0001197 | RLU24197.1 | LOC105284000 | 2.07E-64 |
| Uba3 | FBgn0263697 | RLU24321.1 | LOC105281424 | 0 |

**Table S3.** *O. biroi* Toll pathway gene list. *: genes whose immune function has been questioned by recent studies.

| | Symbol | Flybase ID | O.biroi Homologue CDS ID | O.biroi Homologue Gene ID | BLAST e-value vs D. melanogaster homologue |
|---|---|---|---|---|---|
| Key Components | Tl | FBgn0262473 | RLU17569.1 | LOC105282301 | 0 |
| | tub | FBgn0003882 | RLU17585.1 | LOC105278893 | 1.58E-16 |
| | PGRP-SA | FBgn0030310 | RLU18216.1 | LOC105276273 | 8.03E-52 |
| | spz | FBgn0003495 | RLU20379.1 | LOC105282879 | 1.96E-21 |
| | GNBP3 | FBgn0040321 | RLU20957.1 | LOC105282381 | 5.68E-66 |
| | Myd88 | FBgn0033402 | RLU23134.1 | LOC105286460 | 1.63E-30 |
| | GNBP1 | FBgn0040323 | RLU23175.1 | LOC105281191 | 1.68E-69 |
| | dl | FBgn0260632 | RLU25211.1 | LOC113561388 | 4.37E-148 |
| | cactin | FBgn0031114 | RLU15765.1 | LOC105286140 | 0 |
| Secondary Components | Sumo | FBgn0264922 | RLU17200.1 | LOC105274536 | 1.91E-54 |
| | lwr | FBgn0010602 | RLU17305.1 | LOC105285009 | 1.41E-96 |
| | dx | FBgn0000524 | RLU17824.1 | LOC105283742 | 4.44E-83 |
| | Pitslre | FBgn0016696 | RLU18409.1 | LOC105276119 | 0 |
| | pip | FBgn0003089 | RLU18867.1 | LOC105276110 | 5.26E-180 |
| | mts | FBgn0004177 | RLU19844.1 | LOC105275036 | 0 |
| | tws | FBgn0004889 | RLU19873.1 | LOC105275712 | 0 |
| | Herc4 | FBgn0035207 | RLU20163.1 | LOC105279760 | 0 |
| | Gprk2 | FBgn0261988 | RLU20409.1 | LOC105275024 | 0 |
| | Uba2 | FBgn0029113 | RLU20620.1 | LOC105279706 | 0 |
| | krz | FBgn0040206 | RLU20964.1 | LOC105279882 | 0 |
| | 26-29-p | FBgn0250848 | RLU26281.1 | LOC105284622 | 0 |
| | Pp2A-29B | FBgn0260439 | RLU26725.1 | LOC105278543 | 0 |
| | gro | FBgn0001139 | RLU16486.1 | LOC105274486 | 0 |

| | | | | |
|---|---|---|---|---|
| Pli | FBgn0025574 | RLU17055.1 | LOC105275262 | 0 |
| PGRP-SC2 | FBgn0043575 | RLU17731.1 | LOC105283691 | 1.87E-51 |
| for | FBgn0000721 | RLU19414.1 | LOC105275017 | 0; 0 |
| Spn88Ea | FBgn0028984 | RLU20489.1 | LOC105279932 | 9.96E-74 |
| Doa | FBgn0265998 | RLU21547.1 | LOC105285900 | 0 |
| NT1 | FBgn0261526 | RLU22009.1 | LOC105281847 | 1.50E-52 |
| cact | FBgn0000250 | RLU23038.1 | LOC105287396 | 3.20E-66 |
| Spn42Dd | FBgn0028988 | RLU23116.1 | LOC105285195 | 5.20E-57 |
| senju | FBgn0031676 | RLU23447.1 | LOC105283529 | 2.90E-125 |
| Hrs | FBgn0031450 | RLU23930.1 | LOC105282897 | 4.75E-147 |
| ndl | FBgn0002926 | RLU23967.1 | LOC105283615 | 9.73E-84 |
| Aos1 | FBgn0029512 | RLU24236.1 | LOC105282543 | 2.05E-88 |
| Spn27A | FBgn0028990 | RLU25310.1 | LOC105276796 | 6.81E-83 |
| Traf6 | FBgn0265464 | RLU25883.1 | LOC105280100 | 4.47E-72 |
| gd* | FBgn0000808 | RLU26476.1 | LOC105280274 | 1.55E-69 |

**Table S4.** *O.biroi* additional immune gene list. *: genes whose immune function has been questioned by recent studies.

| Category | Symbol | Flybase ID | O.biroi Homologue CDS ID | O.biroi Homologue Gene ID |
|---|---|---|---|---|
| AMP | waprin 1 | NA | RLU18038.1 | LOC105278357 |
| | waprin 2 | NA | RLU18039.1 | LOC105278356 |
| | SjAPI-like | NA | RLU18089.1 | LOC105278463 |
| | defensin | NA | RLU19863.1 | LOC105274826 |
| | hymenoptaecin | NA | RLU21925.1 | LOC105276730 |
| | apd3 | NA | RLU24908.1 | LOC105275422 |
| | crustin | NA | RLU25224.1 | LOC105277007 |
| | ickin | NA | RLU26638.1 | Ickin1-1 |
| C-type lectin | CTL-101 | NA | RLU15747.1 | LOC105278842 |

| | | | | |
|---|---|---|---|---|
| | CTL5 | NP_001229926.1 | RLU16723.1 | LOC105278676 |
| | CTL-103 | NA | RLU18586.1 | LOC105276307 |
| | CTL12 | XP_006558108.1 | RLU19037.1 | LOC105278999 |
| | CTL1 | XP_001120347.2 | RLU19332.1 | LOC105280740 |
| | CTL-102 | NA | RLU23169.1 | LOC105285211 |
| | CTL3 | XP_006557484.1 | RLU26875.1 | LOC105279962 |
| Cellular response | hemomucin | NA | RLU20915.1 | LOC105282430 |
| | hemolectin | NA | RLU21347.1 | LOC105284524 |
| Dscam* | Dscam2-like-1 | FBgn0265296 | RLU15758.1 | LOC105282206 |
| | Dscam2-like-2 | FBgn0265296 | RLU15763.1 | LOC105282206 |
| | Dscam4-like | FBgn0263219 | RLU16069.1 | LOC105284248 |
| | Dscam2-like-3 | FBgn0265296 | RLU19979.1 | LOC105275687 |
| | Dscam1-like-1 | FBgn0033159 | RLU22744.1 | LOC105287780 |
| | Dscam1-like-2 | FBgn0033159 | RLU22747.1 | LOC105287780 |
| | Dscam3-like | FBgn0033159 | RLU23107.1 | LOC105285240 |
| Galectin | galectin-1 | NA | RLU15819.1 | LOC105285070 |
| | galectin-2 | NA | RLU27311.1 | LOC105280352 |
| JAK-STAT | ken | FBgn0011236 | RLU14795.1 | LOC105286345 |
| | asrij | FBgn0034793 | RLU16763.1 | LOC105274598 |
| | lig | FBgn0020279 | RLU17150.1 | LOC105288194 |
| | E(bx) | FBgn0000541 | RLU18228.1 | LOC105275959 |
| | CycD | FBgn0010315 | RLU18521.1 | LOC105284703 |
| | Ptp61F | FBgn0267487 | RLU19136.1 | LOC105276042 |
| | Socs36E | FBgn0041184 | RLU19257.1 | LOC105280793 |
| | Su(var)2-10 | FBgn0003612 | RLU19689.1 | LOC105287871 |
| | dome | FBgn0043903 | RLU19892.1 | LOC105280778 |
| | apt | FBgn0015903 | RLU20401.1 | LOC105277577 |
| | CycE | FBgn0010382 | RLU21628.1 | LOC105276425 |
| | hop | FBgn0004864 | RLU22133.1 | LOC105276413 |
| | EloB | FBgn0023212 | RLU22753.1 | LOC105287652 |

| | | | | |
|---|---|---|---|---|
| | Cul2 | FBgn0032956 | RLU22941.1 | LOC105285179 |
| | Cul5 | FBgn0039632 | RLU23360.1 | LOC105287145 |
| | Imp | FBgn0285926 | RLU24068.1 | LOC105275815 |
| | mask | FBgn0043884 | RLU24322.1 | LOC105281422 |
| | EloC | FBgn0266711 | RLU24687.1 | LOC105283191 |
| | Cdk2 | FBgn0004107 | RLU24703.1 | LOC105278710 |
| | BRWD3 | FBgn0011785 | RLU24742.1 | LOC105276970 |
| | Stat92E | FBgn0016917 | RLU25734.1 | LOC105281780 |
| | Cnot4 | FBgn0051716 | RLU26990.1 | LOC105278023 |
| | Cdk4 | FBgn0016131 | RLU27343.1 | LOC105280357 |
| | Mkk4 | FBgn0024326 | RLU15198.1 | LOC105283883 |
| | puc | FBgn0243512 | RLU15251.1 | LOC105280616 |
| | pont | FBgn0040078 | RLU16931.1 | LOC105280598 |
| | Uev1A | FBgn0035601 | RLU18291.1 | LOC105276007 |
| | lic | FBgn0261524 | RLU18329.1 | LOC105276227 |
| | CYLD | FBgn0032210 | RLU18404.1 | LOC105276204 |
| | Tak1 | FBgn0026323 | RLU18624.1 | LOC105284644 |
| | grnd | FBgn0032682 | RLU19284.1 | LOC105279843 |
| | ben | FBgn0000173 | RLU20330.1 | LOC105282035 |
| JNK | Tace | FBgn0039734 | RLU20425.1 | LOC105282848 |
| | jra | FBgn0001291 | RLU20611.1 | LOC105277703 |
| | egr | FBgn0033483 | RLU20743.1 | LOC105279798 |
| | hep | FBgn0010303 | RLU21330.1 | LOC105285129 |
| | bsk | FBgn0000229 | RLU21971.1 | LOC105285161 |
| | POSH | FBgn0040294 | RLU23826.1 | LOC105283541 |
| | msn | FBgn0010909 | RLU23835.1 | LOC105282638 |
| | Strip | FBgn0035437 | RLU24195.1 | LOC105285867 |
| | nopo | FBgn0034314 | RLU25130.1 | LOC105277053 |
| | HUWE1 | FBgn0030674 | RLU25305.1 | LOC105276800 |

| | | | | |
|---|---|---|---|---|
| | Traf6 | FBgn0265464 | RLU25883.1 | LOC105280100 |
| | Alg3 | FBgn0011297 | RLU26418.1 | LOC105280429 |
| | Tab2 | FBgn0086358 | RLU27115.1 | LOC105283447 |
| Ig superfamily | immunoglobulin-like domain containing protein 1 | NA | RLU14915.1 | LOC105286610 |
| | immunoglobulin-like domain containing protein 2 | NA | RLU14968.1 | LOC105286893 |
| | immunoglobulin-like domain containing protein 3 | NA | RLU15688.1 | LOC105278806 |
| | immunoglobulin-like domain containing protein 4 | NA | RLU15895.1 | LOC105281448 |
| | immunoglobulin-like domain containing protein 5 | NA | RLU16384.1 | LOC105275896 |
| | immunoglobulin-like domain containing protein 6 | NA | RLU17562.1 | LOC105284967 |
| | immunoglobulin-like domain containing protein 7 | NA | RLU17715.1 | LOC105278392 |
| | immunoglobulin-like domain containing protein 8 | NA | RLU18648.1 | LOC105276208 |
| | immunoglobulin-like domain containing protein 9 | NA | RLU19372.1 | LOC105275527 |
| | immunoglobulin-like domain containing protein 10 | NA | RLU19602.1 | LOC105275771 |
| | immunoglobulin-like domain containing protein 11 | NA | RLU19690.1 | LOC105287868 |
| | immunoglobulin-like domain containing protein 12 | NA | RLU19865.1 | LOC105274831 |

| | | | |
|---|---|---|---|
| immunoglobulin-like domain containing protein 13 | NA | RLU21529.1 | LOC105286657 |
| immunoglobulin-like domain containing protein 14 | NA | RLU22073.1 | LOC105284950 |
| immunoglobulin-like domain containing protein 15 | NA | RLU22074.1 | LOC105286657 |
| immunoglobulin-like domain containing protein 16 | NA | RLU22075.1 | LOC105286657 |
| immunoglobulin-like domain containing protein 17 | NA | RLU22273.1 | LOC105284902 |
| immunoglobulin-like domain containing protein 18 | NA | RLU22480.1 | LOC105279311 |
| immunoglobulin-like domain containing protein 19 | NA | RLU22970.1 | LOC105285222 |
| immunoglobulin-like domain containing protein 20 | NA | RLU23143.1 | LOC105275172 |
| immunoglobulin-like domain containing protein 21 | NA | RLU23270.1 | LOC105280421 |
| immunoglobulin-like domain containing protein 22 | NA | RLU23697.1 | LOC105283797 |
| immunoglobulin-like domain containing protein 23 | NA | RLU23714.1 | LOC105283839 |
| immunoglobulin-like domain containing protein 24 | NA | RLU23791.1 | LOC105280531 |
| immunoglobulin-like domain containing protein 25 | NA | RLU24039.1 | LOC105283549 |
| immunoglobulin-like | NA | RLU24521.1 | LOC105274676 |

| | | | |
|---|---|---|---|
| domain containing protein 26 | | | |
| immunoglobulin-like domain containing protein 27 | NA | RLU24527.1 | LOC105287749 |
| immunoglobulin-like domain containing protein 28 | NA | RLU24528.1 | LOC105288120 |
| immunoglobulin-like domain containing protein 29 | NA | RLU24552.1 | LOC105276843 |
| immunoglobulin-like domain containing protein 30 | NA | RLU24621.1 | LOC105277142 |
| immunoglobulin-like domain containing protein 31 | NA | RLU24780.1 | LOC105277040 |
| immunoglobulin-like domain containing protein 32 | NA | RLU25022.1 | LOC105277484 |
| immunoglobulin-like domain containing protein 33 | NA | RLU25075.1 | LOC105278694 |
| immunoglobulin-like domain containing protein 34 | NA | RLU25403.1 | LOC105282923 |
| immunoglobulin-like domain containing protein 35 | NA | RLU25435.1 | LOC105287160 |
| immunoglobulin-like domain containing protein 36 | NA | RLU25619.1 | LOC105284017 |
| immunoglobulin-like domain containing protein 37 | NA | RLU26157.1 | LOC105282464 |
| immunoglobulin-like domain containing protein 38 | NA | RLU26631.1 | LOC105274945 |
| immunoglobulin-like domain containing protein | NA | RLU26636.1 | LOC105274944 |

| | | | | |
|---|---|---|---|---|
| | 39 | | | |
| | immunoglobulin-like domain containing protein 40 | NA | RLU27434.1 | LOC105278529 |
| **Lysozyme** | Lys_a | NA | RLU26681.1 | LOC113562164 |
| | Lys_b | NA | RLU27337.1 | LOC105285451 |
| **PGRP** | PGRP-SD | FBgn0035806 | RLU16632.1 | LOC105274606 |
| | PGRP-SC2 | FBgn0043575 | RLU17731.1 | LOC105283691 |
| | PGRP-LB | FBgn0037906 | RLU17733.1 | LOC105283690 |
| | PGRP-LC | FBgn0035976 | RLU18495.1 | LOC105279031 |
| | PGRP | NA | RLU18496.1 | LOC105279033 |
| **PPO** | PPO | NA | RLU15745.1 | LOC105278857 |
| | PPO | NA | RLU24656.1 | LOC105277104 |
| **RNAi** | Ago2 | FBgn0087035 | RLU15437.1 | LOC105284026 |
| | Dcr2 | FBgn0034246 | RLU23508.1 | LOC105285125 |
| | Dcr1 | FBgn0039016 | RLU26333.1 | LOC105283491 |
| **Scavenger receptor** | Sr-CII | FBgn0020377 | RLU15813.1 | LOC105277762 |
| | NimX | NA | RLU18652.1 | LOC113562625 |
| | eater | FBgn0243514 | RLU18653.1 | LOC105285560 |
| | NimA | FBgn0261514 | RLU18661.1 | LOC105285553 |
| | Snmp1 | FBgn0260004 | RLU19313.1 | LOC105280335 |
| | CG10345 | FBgn0027562 | RLU19322.1 | LOC105279695 |
| | emp-1 | FBgn0010435 | RLU19684.1 | LOC105287875 |
| | emp-2 | FBgn0010435 | RLU19685.1 | LOC105287876 |
| | CG40006 | FBgn0058006 | RLU19750.1 | LOC105285836 |
| | Croquemort | FBgn0015924 | RLU20261.1 | LOC105282838 |
| | Sr-CII | FBgn0020377 | RLU21276.1 | LOC105276575 |
| | dsb | FBgn0035290 | RLU21929.1 | LOC105276733 |
| | santa maria | FBgn0025697 | RLU25059.1 | LOC105287562 |
| | Draper-like | FBgn0027594 | RLU27105.1 | LOC105278058 |
| **Serine protease** | ObirSP17 | NA | RLU16061.1 | LOC105278864 |

| | | | |
|---|---|---|---|
| ObirSP20 | NA | RLU16063.1 | LOC105278891 |
| ObirSP39 | NA | RLU16222.1 | LOC105286116 |
| ObirSP32 | NA | RLU16646.1 | LOC105288169 |
| ObirSP64 | NA | RLU17773.1 | LOC105288146 |
| ObirSP43 | NA | RLU18447.1 | LOC105281621 |
| ObirSP65 | NA | RLU18655.1 | LOC105279599 |
| ObirSP47 | NA | RLU18662.1 | LOC105279641 |
| ObirSP26 | NA | RLU19090.1 | LOC105284678 |
| ObirSP13 | NA | RLU19126.1 | LOC105284666 |
| ObirSP12 | NA | RLU19218.1 | LOC105287866 |
| ObirSP14 | NA | RLU19600.1 | LOC105275667 |
| ObirSP10 | NA | RLU19726.1 | LOC105279727 |
| ObirSP38 | NA | RLU19794.1 | LOC105275537 |
| ObirSP6 | NA | RLU20046.1 | LOC105279292 |
| ObirSP22 | NA | RLU20199.1 | LOC105286577 |
| ObirSP1 | NA | RLU20303.1 | LOC105287672 |
| ObirSP50 | NA | RLU20632.1 | LOC105279732 |
| ObirSP44 | NA | RLU20641.1 | LOC105286989 |
| ObirSP18 | NA | RLU21016.1 | LOC105279854 |
| ObirSP56 | NA | RLU21041.1 | LOC105282413 |
| ObirSP57 | NA | RLU21223.1 | LOC105276255 |
| ObirSP34 | NA | RLU21281.1 | LOC105276509 |
| ObirSP63 | NA | RLU21377.1 | LOC105276054 |
| ObirSP7 | NA | RLU21684.1 | LOC105276484 |
| ObirSP9 | NA | RLU21689.1 | LOC105276484 |
| ObirSP60 | NA | RLU22110.1 | LOC105284336 |
| ObirSP4 | NA | RLU22189.1 | LOC105276676 |
| ObirSP19 | NA | RLU22623.1 | LOC105281252 |
| ObirSP25 | NA | RLU22796.1 | LOC105281021 |

| | | | |
|---|---|---|---|
| ObirSP52 | NA | RLU22998.1 | LOC105286473 |
| ObirSP15 | NA | RLU23132.1 | LOC105286466 |
| ObirSP5 | NA | RLU23266.1 | LOC105283841 |
| ObirSP23 | NA | RLU23419.1 | LOC105282443 |
| ObirSP31 | NA | RLU23675.1 | LOC105280626 |
| ObirSP33 | NA | RLU24131.1 | LOC105284681 |
| ObirSP35 | NA | RLU24422.1 | LOC105281367 |
| ObirSP29 | NA | RLU24611.1 | LOC105285724 |
| ObirSP55 | NA | RLU24696.1 | LOC105277126 |
| ObirSP2 | NA | RLU25637.1 | LOC105281574 |
| ObirSP11 | NA | RLU25863.1 | LOC105281659 |
| ObirSP61 | NA | RLU25864.1 | LOC105281656 |
| ObirSP48 | NA | RLU26278.1 | LOC105283349 |
| ObirSP27 | NA | RLU26344.1 | LOC105281464 |
| ObirSP45 | NA | RLU26369.1 | LOC105278064 |
| ObirSP21 | NA | RLU26476.1 | LOC105280274 |
| ObirSP16 | NA | RLU26607.1 | LOC105286745 |
| ObirSP37 | NA | RLU26608.1 | LOC105286748 |
| ObirSP40 | NA | RLU26774.1 | LOC105278563 |
| ObirSP41 | NA | RLU26777.1 | LOC105278563 |
| ObirSP8 | NA | RLU26778.1 | LOC105278512 |
| ObirSP49 | NA | RLU26811.1 | LOC105281465 |
| ObirSP3 | NA | RLU26838.1 | LOC105278506 |
| ObirSP59 | NA | RLU27153.1 | LOC105281477 |
| ObirSP28 | NA | RLU27155.1 | LOC105281472 |
| ObirSP36 | NA | RLU27156.1 | LOC105281472 |
| ObirSP62 | NA | RLU27227.1 | LOC105286748 |
| ObirSP30 | NA | RLU27228.1 | LOC105286745 |
| ObirSP42 | NA | RLU27333.1 | LOC105285446 |

| | | | | |
|---|---|---|---|---|
| | ObirSP51 | NA | RLU27368.1 | LOC105278569 |
| | ObirSP54 | NA | RLU27374.1 | LOC105278548 |
| | ObirSP24 | NA | RLU27383.1 | LOC113562287 |
| | ObirSP58 | NA | RLU27477.1 | LOC105280438 |
| | ObirSP53 | NA | RLU27480.1 | LOC109611042 |
| | ObirSP46 | NA | RLU27482.1 | LOC105280485 |
| STING | IKKŒµ | FBgn0086657 | RLU16469.1 | LOC105288233 |
| | Sting | FBgn0033453 | RLU21634.1 | LOC105276704 |
| | cGlr1 | FBgn0034047 | RLU23970.1 | LOC105279211 |
| TEP | TEP_c | NA | RLU16697.1 | LOC105274538 |
| | TEP_b | NA | RLU17945.1 | LOC113562813 |
| | TEP_a | NA | RLU17946.1 | LOC105282295 |
| | TEP_d | NA | RLU23037.1 | LOC105287371 |
| TLR | Toll-6 | NA | RLU16709.1 | LOC105282942 |
| | Toll-unclear (2/7?) | NA | RLU16722.1 | LOC105275904 |
| | Toll-8 | NA | RLU16738.1 | LOC105282937 |
| | Toll-unclear (2/7?) | NA | RLU24395.1 | LOC105276942 |

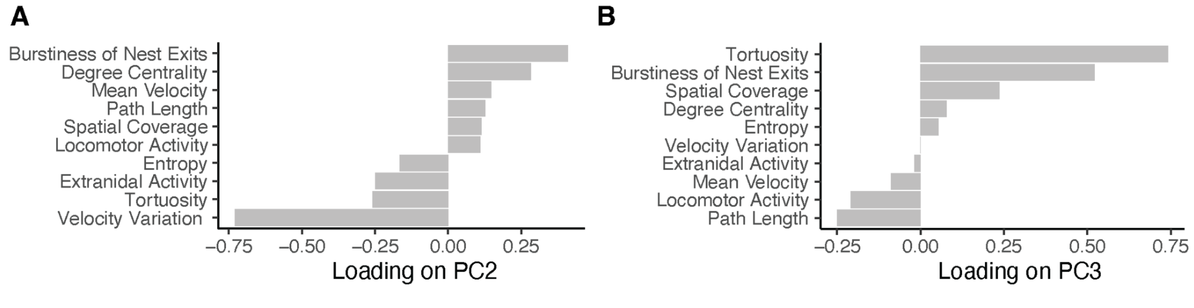


**Figure S1.** Loadings of behavioural metrics on (A) PC2 and (B) PC3 in the experiment investigating the association between individual behaviour and immune-related gene expression.

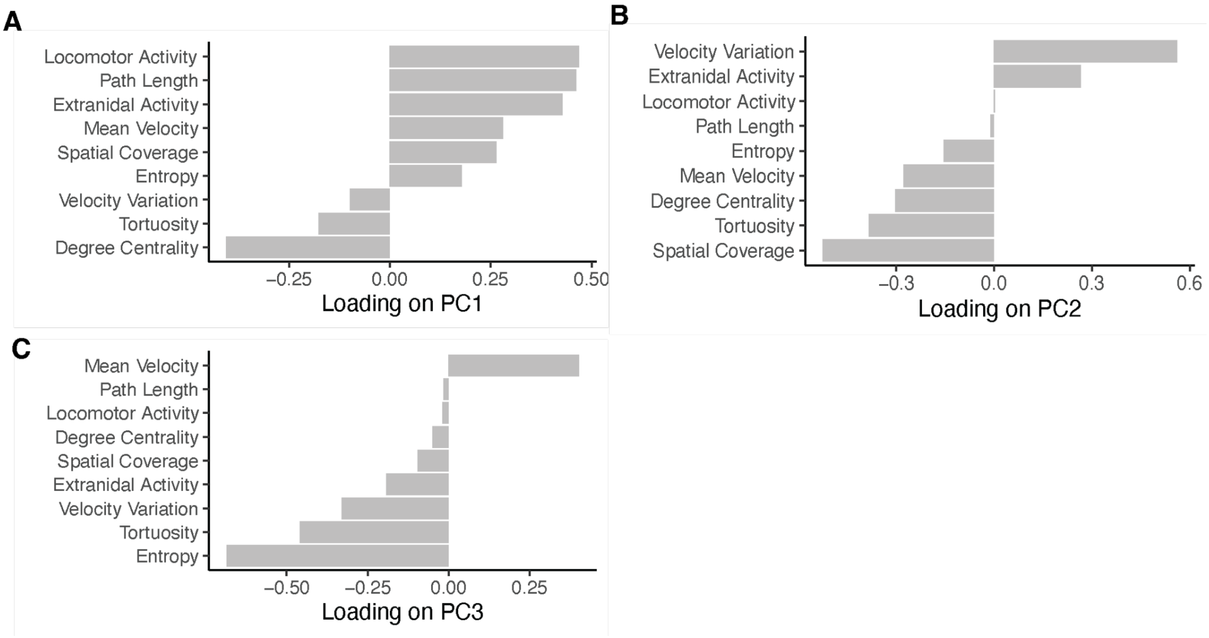


**Figure S2.** Loadings of behavioural metrics on (A) PC1, (B) PC2 and (C) PC3 in the experiment investigating the association between individual behaviour and survival upon fungal exposure.

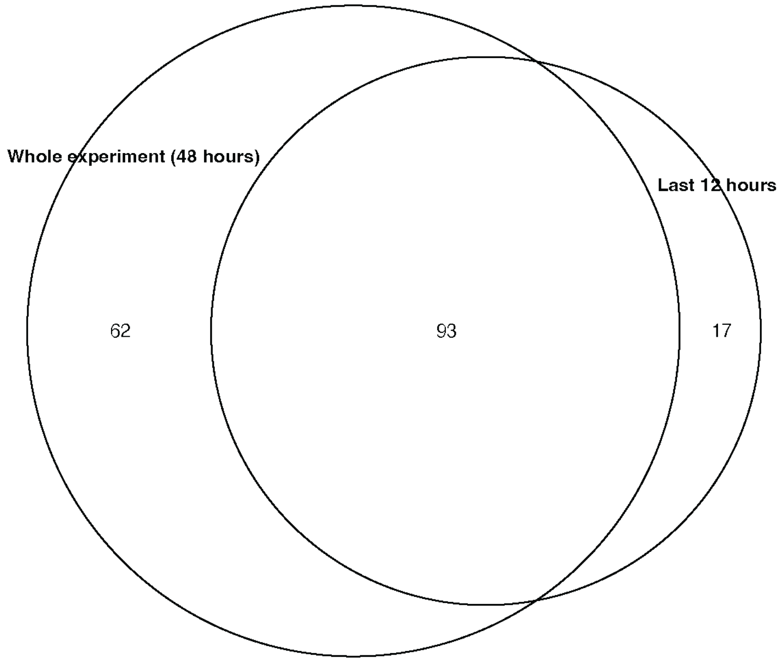


**Figure S3.** Overlap of differentially expressed genes based on behavioural data of the whole experiment (48 hours) vs. the last 12 hours.
